# Mouse models of immune dysfunction: Their neuroanatomical differences reflect their anxiety-behavioural phenotype

**DOI:** 10.1101/2021.06.14.448406

**Authors:** Darren J. Fernandes, Shoshana Spring, Christina Corre, Andrew Tu, Lily R. Qiu, Christopher Hammill, Dulcie A. Vousden, T. Leigh Spencer Noakes, Brian J. Nieman, Dawn M.E Bowdish, Jane A. Foster, Mark R. Palmert, Jason P. Lerch

**Affiliations:** Mouse Imaging Centre, The Hospital for Sick Children, Toronto, ON, Canada; Department of Medical Biophysics, The University of Toronto, Toronto, ON, Canada; Department of Neurosciences and Mental Health, The Hospital for Sick Children, Toronto, ON, Canada; Division of Endocrinology, The Hospital for Sick Children, Toronto, ON, Canada; Department of Pathology and Molecular Medicine, McMaster University, Hamilton, ON, Canada; Department of Psychiatry and Behavioral Neurosciences, McMaster University, Hamilton, ON, Canada; Departments of Paediatrics and Physiology, The University of Toronto, Toronto, ON, Canada; Department of Preclinical Imaging, Wellcome Centre for Integrative Neuroimaging, University of Oxford, Oxford, United Kingdom; Translational Medicine, The Hospital for Sick Children, Toronto, ON, Canada; Ontario Institute for Cancer Research, Toronto, ON, Canada

**Author notes:** First authors who contributed equally (Darren J. Fernandes, Shoshana Spring). Senior authors who contributed equally (Mark R. Palmert, Jason P. Lerch).

**Keywords:** MRI, immune system, mouse brain, behaviour, anxiety

## Abstract

Extensive evidence supports the role of the immune system in modulating brain function and behaviour. However, past studies have revealed striking heterogeneity in behavioural phenotypes produced from immune system dysfunction. Using magnetic resonance imaging (MRI), we studied the neuroanatomical differences among 11 distinct genetically-modified mouse lines (n=371), each deficient in a different element of the immune system. We found a significant and heterogeneous effect of immune dysfunction on the brains of both male and female mice. However, by imaging the whole brain and using bayesian hierarchical modelling, we were able to identify patterns within the heterogeneous phenotype. Certain structures -- such as the corpus callosum, midbrain, and thalamus -- were more likely to be affected by immune dysfunction. A notable brain-behaviour relationship was identified with neuroanatomy endophenotypes across mouse models clustering according to anxiety-like behaviour phenotypes reported in literature, such as altered volume in brains regions associated with promoting fear response (e.g., the lateral septum and cerebellum). Interestingly, genes with preferential spatial expression in the most commonly affected regions are also associated with multiple sclerosis and other immune-mediated diseases. In total, our data suggest that the immune system modulates anxiety behaviour through well-established brain networks.

## Introduction

The immune system plays an important role in brain-behaviour interactions. Immune dysregulation within the maternal-fetal environment is associated with an increased risk of central nervous system (CNS) disorders[1–3]. In particular, maternal illness during pregnancy has been linked with increased rates of impaired neurodevelopment and disorders such as autism spectrum disorder[4], schizophrenia[5], and multiple sclerosis[3]. Several cytokines have been shown to play a role in mediating the neurobehavioral consequences of infection including interleukin (IL)-6, −17, and −2[6–8]. Given that immune disruption in early life can produce long-term changes in neuronal function, it is important to study how the immune system influences the brain.

Mice have been used to investigate how the immune system modulates brain structure and behaviour. Altering expression of genes involved with production of cytokines (such as IL-6[9], IL-10[10], and IL-18[11]) and cell-mediated adaptive immune response (such as Ighm[12], CD4[13], CD8[14], Rag1[15], and Rag2[16]) is associated with changes in anxiety behaviours. The modulation of anxiety is also observed when altering other components of the immune system. Male Nos2 mutant mice showed a statistically significant anxiety-like phenotype in the elevated plus maze (EPM)[17]. Kit male mutants, which are mast cell-deficient, have a greater anxiety-like phenotype in both OFT and EPM[18].

Immune system effects on the brain and, ultimately, behaviour are heterogenous and multifaceted. Existing studies generally examine immune system activations/mutations and brain regions or behaviours in isolation, thereby sacrificing the ability to observe broader patterns that may explain the observed heterogeneity. Structural magnetic resonance imaging (MRI) is a useful methodology for studying relationships between genetics, brain regions, and behaviour with several studies demonstrating a strong link between the volume of brain structures and organism behaviour[19, 20]. As structural MRI has the benefit of whole brain coverage without the constraint to identify regions-of-interest ahead of time, it allows identification of novel, unanticipated phenotypes in mouse models of human disease. For example, we previously showed using *ex vivo* MRI that mice lacking all functional T cells demonstrated a loss of sexual dimorphism in many brain regions[12]. We corroborated this finding with a loss of sexual differentiation in activity-related behaviors, highlighting the importance of T cells for the development of sex differences in neuroanatomy and behavior. However, such studies remain rare and the brain effects of alterations in other elements of the immune system are unknown. High-throughput data acquisition and analysis of structural MRI allows for investigation of similarities and differences across a range of immune system mutations.

Thus, here we used whole brain MRI to explore the effect of immune system mutations on the brains of male and female mice. We imaged the brains of 11 genetically-engineered mouse lines, each deficient in different elements of the immune system, using a standard pipeline for specimen preparation, data acquisition, and image registration. Brain structure was highly susceptible to immune dysfunction and we found regional variations in this susceptibility. The most susceptible regions tended to express genes associated with immune-mediated diseases. We also identified that strains with similar anxiety behavioural phenotypes displayed analogous neuroanatomical effects. Lastly, we identified candidate brain regions and networks that may be involved in modulating the anxiety behaviours seen in mouse models of immune dysfunction.

## Results

### Immune dysfunction affects male and female neuroanatomy

A significant effect of strain was observed on the volumes of nearly all brain structures (Figure 1A) and this effect was similar for male and female mice (Supplementary Figure 6). While the effect of strain was highly significant, the magnitude and direction of these effects showed a great deal of heterogeneity (Figure 1B for females, Supplementary Figure 5 for males, Supplementary Table 4 for complete list). For example, mutants lacking cytokines IL-6, IL-10, and IL-18 had a similar phenotype of larger cerebellum compared to wild-type. However, they differed in phenotypes of the left frontal association cortex with IL-10 being smaller than wild-type, and IL-6 and IL-18 being larger.

**Figure 1:**
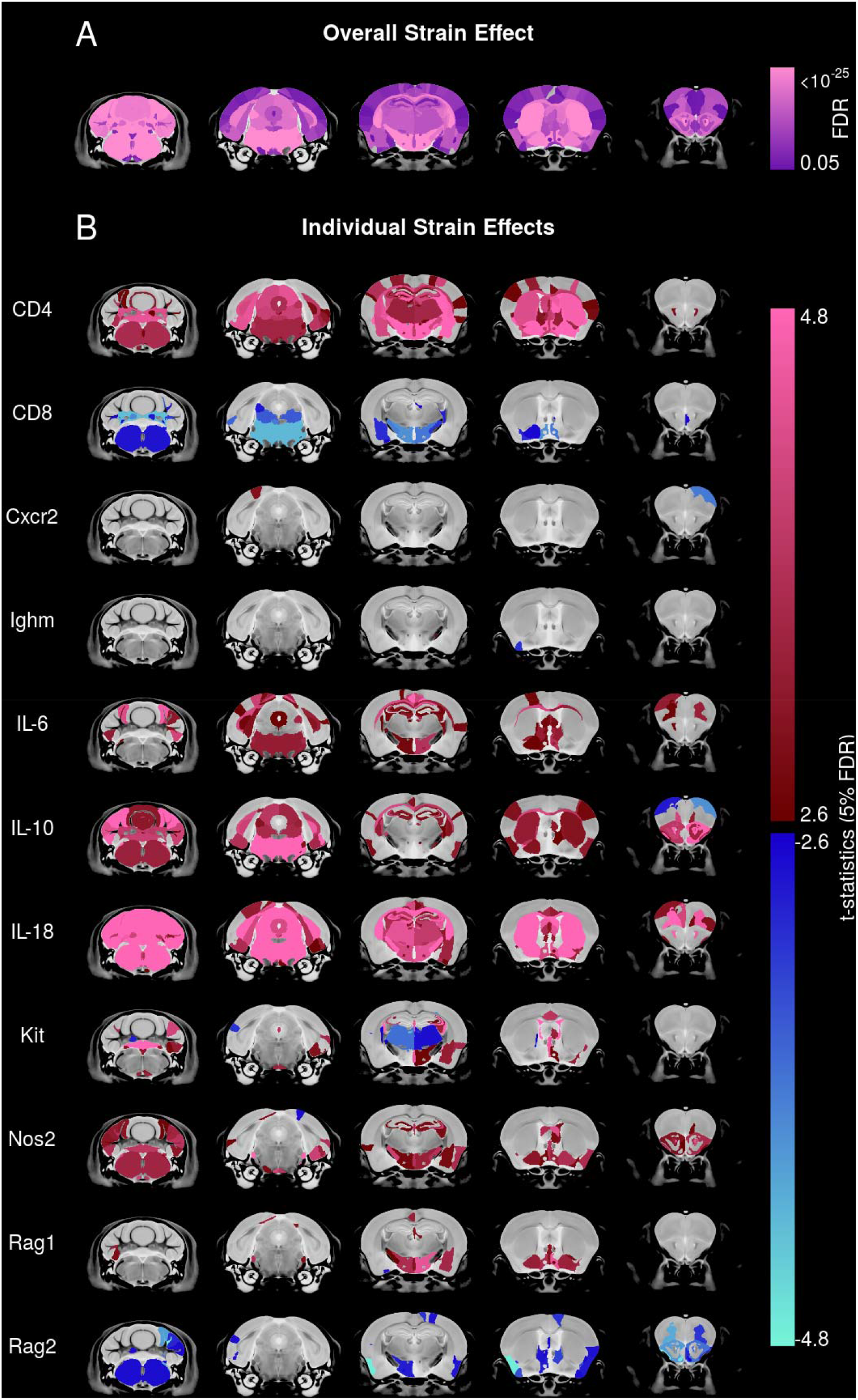
Immune system mutations have a highly heterogeneous effect on mouse brain anatomy. (A) Nearly all brain structures showed a significant effect of strain evaluated using F-statistics from ANOVA. (B) The directional effect in females of the various mutant strains relative to the wild-type strains is visualized using t-statistics and shows a heterogeneous neuroanatomical phenotype. Regions larger or smaller in mutants relative to wild-type are given maroon-pink and blue-turquoise colours, respectively, if effects are <5%FDR. Saturated colours represent effects <0.01% FDR.

To find important patterns through these heterogeneous effects, a bayesian hierarchical model (BHM) was fitted to the neuroanatomy data. Consistent with the frequentist analysis (Figure 1A), all brain structures had a high probability of having moderate effect-size magnitudes (95% credible interval exceeded 0.50 for all structures). However, certain structures were particularly sensitive to strain and had a high chance of having large effect-sizes (Figure 2A). For example, the Midbrain (Figure 2B), Corpus callosum (Figure 2C), Dorsal striatum (Figure 2D), and Thalamus (Figure 2E) all had among the highest probabilities of having large effect-sizes across the 11 strains evaluated.

**Figure 2:**
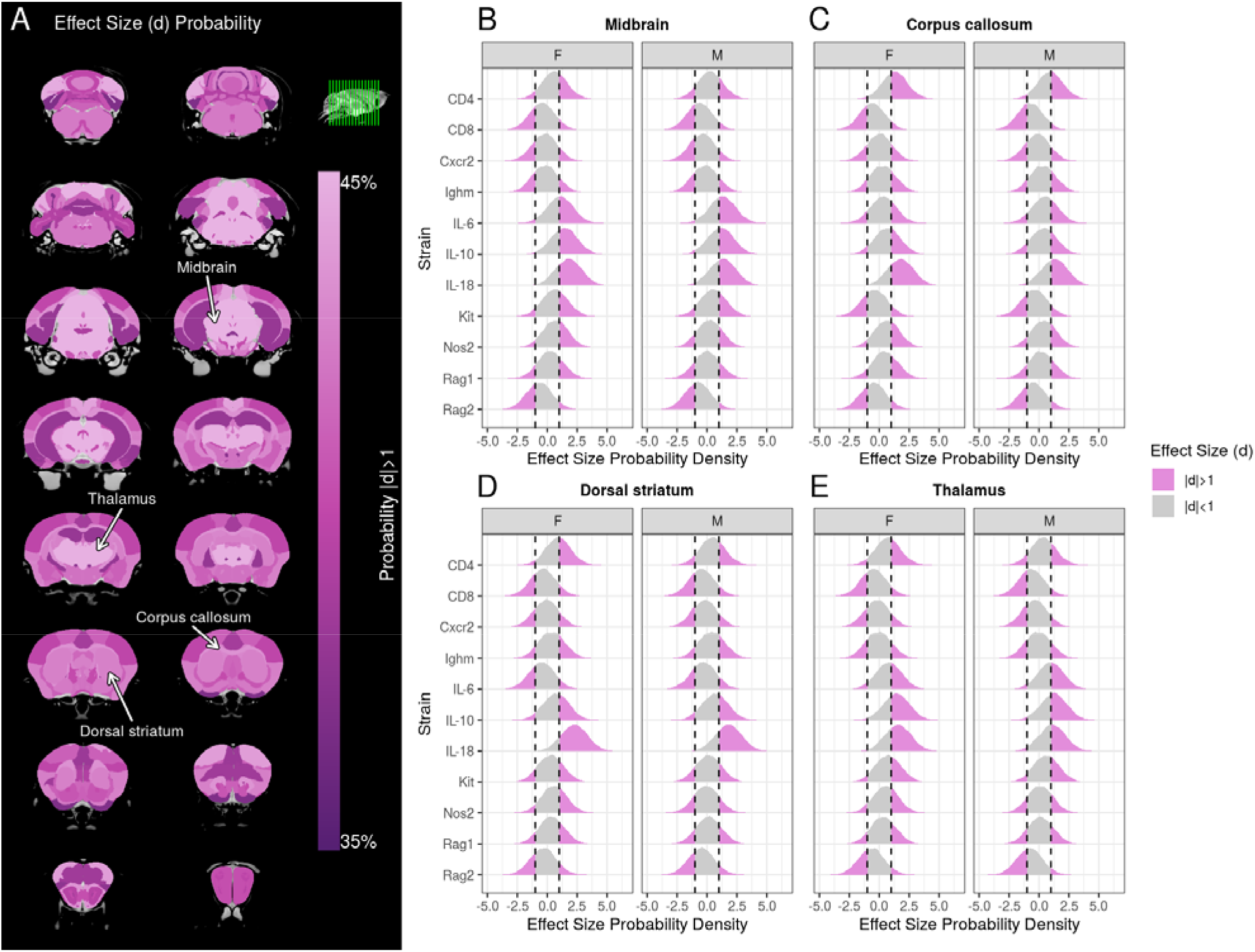
Brain regions showed variations in susceptibility to volume changes due to immune system mutations. (A) The probability of a brain region having a large effect-size magnitude in mutant strains. Probability distributions of effect-sizes (x-axis) for various structures -- (B) Midbrain, (C) Corpus callosum, (D) Dorsal striatum, (E) Thalamus -- across the different mutant strain (y-axis), for both female (F) and male (M) mice. Vertical dashed lines represent effect sizes of ±1, and probability distribution within this interval is shaded grey. Integrating the area of the probability density outside this interval (pink) provides the probability that immune system mutations result in a large effect-size magnitude. These four structures had the highest probability of large effect-size magnitude.

### Neuroanatomical differences among strains cluster by anxiety phenotype

Given immune strains had heterogeneous neuroanatomical phenotypes, we investigated whether clustering these phenotypes could reveal important underlying biology. Because prior literature has shown that immune deregulation causes heterogeneous anxiety-like behaviours (summarized in Supplementary Table 3), we assessed whether strains with a similar neuroanatomical endophenotype clustered by anxiety behaviour. We found that mutant strains with similar anxiety behavioural phenotypes had similar neuroanatomical endophenotypes (Figure 3A inset). This association is particularly driven by strains with increased anxiety behaviours, as visualised in a network plot (Figure 3A). We then assessed which brain structures in particular drive this association with anxiety behaviour. Nearly every structure had at least a moderate effect-size of anxiety (Figure 3B); the culmen and lateral septum were plotted as representative structures (Figure 3C). As the Cxcr2 mutant strain did not have a documented anxiety phenotype, we assumed this strain had no anxiety-like behaviours for this analysis; repeating this analysis excluding this strain resulted in similarly strong grouping by anxiety phenotype.

**Figure 3:**
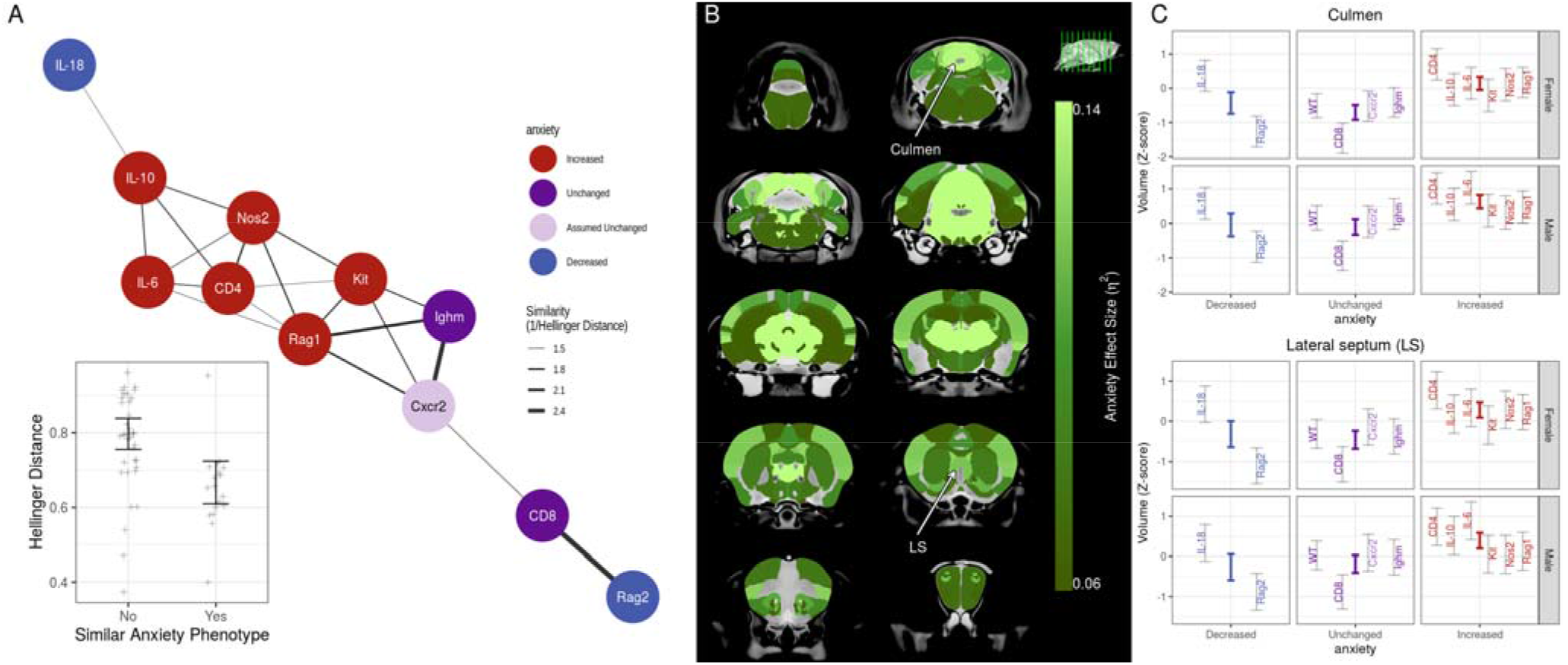
Mutant strains with similar anxiety behavioural phenotype have similar neuroanatomy endophenotypes. For each pair of strains, dissimilarity of endophenotypes was assessed using Hellinger distance and visualized using a network (thicker edges imply greater similarity). (A) Strains with increased anxiety behaviours (red nodes) had similar endophenotypes and clustered together in the network. A similar pattern was seen for unchanged anxiety behaviours (dark purple), but not decreased anxiety behaviour (blue). Cxcr2 anxiety phenotype is not known and assumed unchanged (light purple). The inset plot shows that pairs of strains (represented as crosses) with similar anxiety phenotype had similar neuroanatomy endophenotypes (p<0.01 from permutation testing). (B) The effect size (*η*^2^) for the anxiety grouping was computed for each structure. The green colour bar represents at least medium effect sizes and saturates for large effect sizes. (C) The Culmen and Lateral septum were chosen as representative examples to illustrate large effect-size for anxiety grouping. The 95%-credible-interval of predicted volume for each strain (gray bars) and anxiety phenotype (coloured bars) are shown.

### Affected neuroanatomy has preferential spatial expression of genes associated with immune-mediated disease

We sought to link the neuroanatomical findings with immune dysfunction by comparison with gene expression patterns from the ABI gene expression atlas<u>[41]</u>. For this purpose, a region-of-interest (ROI) was defined from the MRI results by selecting the set of 25 structures with the highest median effect-size magnitude (shown in Figure 4C first row). Spatial gene expression analysis<u>[43, 44]</u> was used to identify genes that were preferentially expressed within this ROI using a fold-change measure (average expression in ROI divided by average expression in the brain). Using the DisGeNet<u>[46]</u> and NCBI<u>[45]</u> databases, we then identified diseases in which these preferentially expressed genes have been implicated. Many of the disease terms with significant enrichment have known or suspected immune-mediated pathophysiology; such as Parkinson Disease (p<10^−8^,FDR<10^−4^), Alzheimer’s disease (p<10^−8^,FDR<10^−4^), and multiple sclerosis (MS) (p<10^−4^,FDR=0.016).

**Figure 4:**
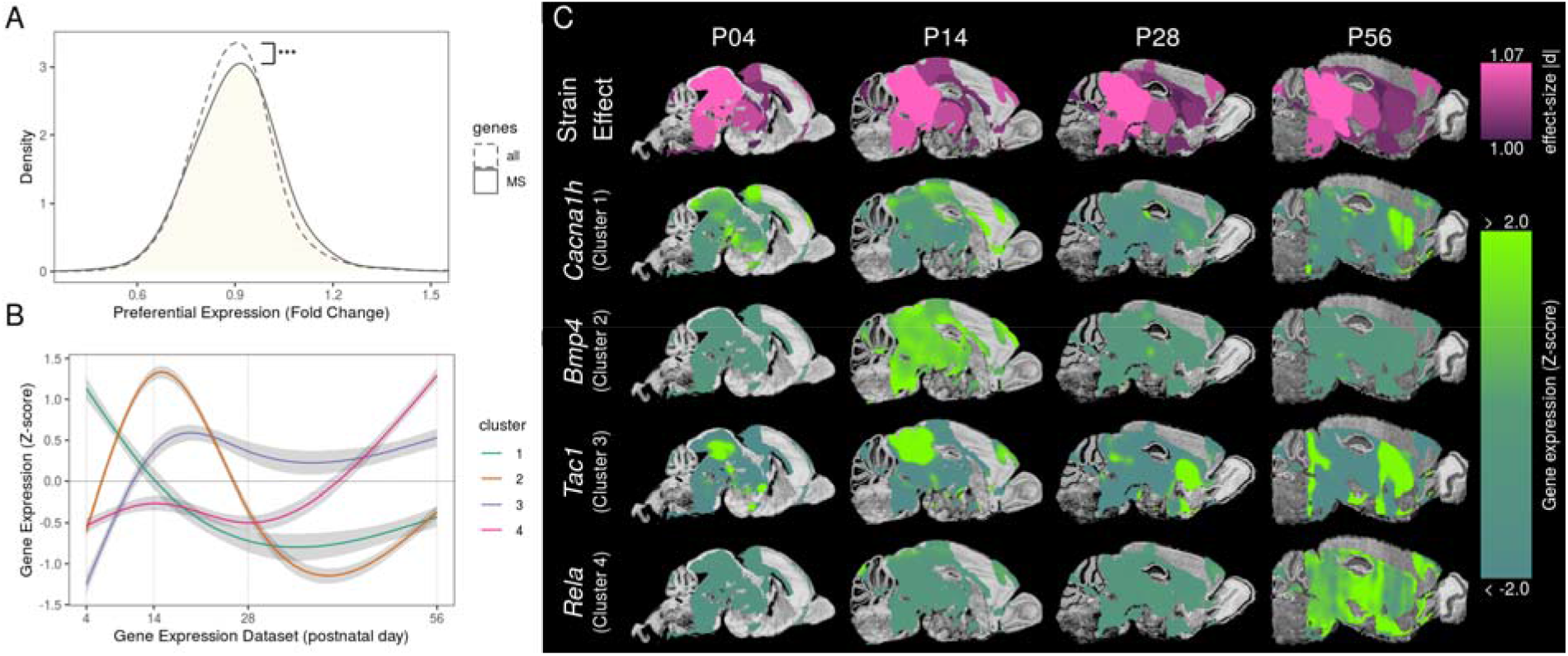
Brain regions susceptible to immune system mutations have preferential spatial expression of genes involved with multiple sclerosis (MS). The top 25 brain structures with highest effect-size magnitudes are shown in C (first row) and constitute the region-of-interest (ROI). For all genes in the mouse genome, preferential spatial expression was assessed using a fold-change measure (i.e. gene expression signal in ROI relative to the whole brain). (A) Genes associated with MS (solid line) had significantly higher expression in the ROI compared to the genome (two-sided KS test D=0.05,p<0.10^−3^). (B) MS genes showed 4 different clusters of temporal expression within the ROI. Shaded region represents the 95% credible interval. (C) A representative example from each cluster was chosen to visualize gene expression signals within the ROI over the course of neurodevelopment. Each example was closest to its respective cluster’s centroid.

As there is strong evidence that MS is an immune-mediated disease<u>[49]</u>, we decided to explore it further. Consistent with the previous enrichment analysis, genes associated with MS tended to have higher expression in brain regions sensitive to immune dysfunction (Figure 4A). We also clustered the expression patterns of MS-associated genes in the ROI through the course of postnatal development<u>[47]</u> and found 4 main clusters of genes. Two clusters had low expression in adulthood with expression peaks in early life: Cluster 1 had a peak at ~P4 and Cluster 2 peaked later ~P14. The other two clusters had high expression in adulthood: Cluster 3 had high expression starting from ~P14 and Cluster 4 starting from ~P56 onward.

## Discussion

Using neuroanatomical imaging with MRI, we confirm significant and widespread effects of the immune system on the mouse brain using a selection of immune-related genetically-engineered mouse lines. Our results are consistent with other studies. For example, Rag1 mutants did not have a gross brain morphology phenotype detectable in 5μm coronal sections taken through the dorsal hippocampus[15]. Our analysis also showed no significantly altered neuroanatomy in the dorsal hippocampus of Rag1 mutants. We found thalamic changes in mast cell-deficient Kit mutants, which is consistent with mast cells migrating along blood vessels of the hippocampus and fimbria during postnatal development and penetrating into the thalamus, where they remain throughout adulthood[50].

Analysis of the heterogeneous structural findings for association with behavioural phenotypes revealed important insights and suggests that neuroanatomical endophenotypes cluster strongly with anxiety-like behavioural phenotypes. Previous use of mouse models has revealed several neural circuits associated with producing anxiety behaviours[51, 52], but it is not known which of these neural circuits are responsible for driving anxiety behaviours in models of immune dysfunction. We found that alterations in structure of the lateral septum and hypothalamus cluster with anxiety. These brain regions are part of the septohypothalamic circuit, whose activation can promote persistent anxiety[53]. We also found that changes in parts of the cerebellum and the midbrain were correlated with anxiety. The cerebellum is connected to many brain regions and there is emerging evidence that it is an important link between the fear- and anxiety-related brain networks[54]. In particular, its connection to the midbrain has been shown to be essential for unconditioned freezing behaviour in rodents[55]. Further investigation of these brain networks may prove useful in understanding how the immune system affects anxiety behaviours. Interestingly, anxiety disorders are a notable comorbid condition in neurological disorders associated with immune dysfunction. For example, anxiety disorders are three times more prevalent in MS patients than in the general population[56]. The prevalence of anxiety disorders in Parkinsons’ is 31%[57].

Our analysis of anxiety was limited by the requirement that we treat anxiety behaviours as ordinal levels, rather than as a spectrum, as the current available literature does not consistently employ the same behavioural assays or report sex differences. Nonetheless, we believe we have identified an important link between immune system dysregulation and anxiety, and hope future studies can gather more consistent behavioural data for both sexes. Initiatives like the International Mouse Phenotyping Consortium (IMPC) could be tremendously useful as they would have consistent behavioural assays for every single-gene mutation in the mouse. This would allow future studies to not only investigate the association between immune mutations and anxiety, but other behaviours as well.

Immune dysfunction plays an important role in many neurological disorders, but the impact of the immune system is difficult to study because the etiology is highly polygenic. For example, MS is associated with over 1000 genetic risk variants[46], but many of these variants are found in proximity to genes with known immune function (ex. major histocompatibility complex on chromosome 6p21.3)[58]. Although requiring further validation, our data provide a potential new avenue for examining the basis of these interactions. By studying neuroanatomy of several mouse models, our methodology may be useful in finding patterns underlying polygenic immune dysfunction. The corpus callosum, thalamus, striatum, and midbrain were amongst the most-affected structures across all mouse models, which reflects neurological findings in autoimmune disorders[59–62]. While there were many mouse genes preferentially expressed in these brain regions, several of these genes are homologous to human genes associated with MS. This finding suggests that expression patterns of MS genes could be followed through neurodevelopment to identify when their expression peaks, pointing to possible developmental time-windows for further study. In addition to MS, other diseases suspected to be immune-mediated also showed enrichment in preferentially-expressed genes, such as Parkinson’s[63] and Alzheimer’s disease[64]. Our data supports the hypothesis that, while there are many genetic variants associated with immune system dysregulation, certain brain regions may be more sensitive to alterations in the immune system. Thus, further study of mouse models with targeted immune system mutations could uncover potential insights regarding neurological symptoms in polygenic immune-mediated disorders.

In summary, we imaged the neuroanatomy of 11 different mouse mutants, each deficient in particular components of the immune system. By using a consistent high-throughput protocol for data acquisition and analysis, we characterized the diverse effects of immune dysregulation on the brain and identified patterns underlying this heterogeneity. The observed neuroanatomical differences in the mouse models of immune dysfunction are clustered by their anxiety phenotype, recapitulating known brain networks implicated in modulating anxious behaviours. Interestingly, exploratory analysis of gene expression shows that the brain regions that are most affected by immune dysfunction have preferential spatial expression of genes associated with MS and other immune-mediated diseases.

## Materials and Methods

### Mice

The study consisted of 371 mice (Table 1) from 11 different genetically-engineered strains obtained from Jackson Labs (JAX, Bar Harbor, ME). These strains were chosen as they encapsulate the different components of the immune system (Supplementary Figure 1), thereby letting us study the effect of these components on brain anatomy. Homozygous mutant mice were used for study purposes. All breeding colonies were maintained using the homozygous x homozygous mutant mating scheme with two exceptions: Cxcr2 colony was maintained using a heterozygous x heterozygous scheme due to the fragility of homozygotes<u>[21]</u>, and Rag2 experimental mice were ordered directly from JAX. Genotypes were determined from tail biopsies using real-time PCR (probes available from Transnetyx, Cordova, USA or Dr. J. Foster, McMaster University, Hamilton, Canada) or fur colour in the case of the Kit strain. The control group consisted of C57Bl/6J mice and Cxcr2 wild-type littermates<u>[22]</u>. On P65±3 (mean±max range), mice were sacrificed for fixation. The weight, nose-to-rump length, and post-fixation organ weight are reported in Supplementary Table 1 and Supplementary Figure 2.

**Table 1:**
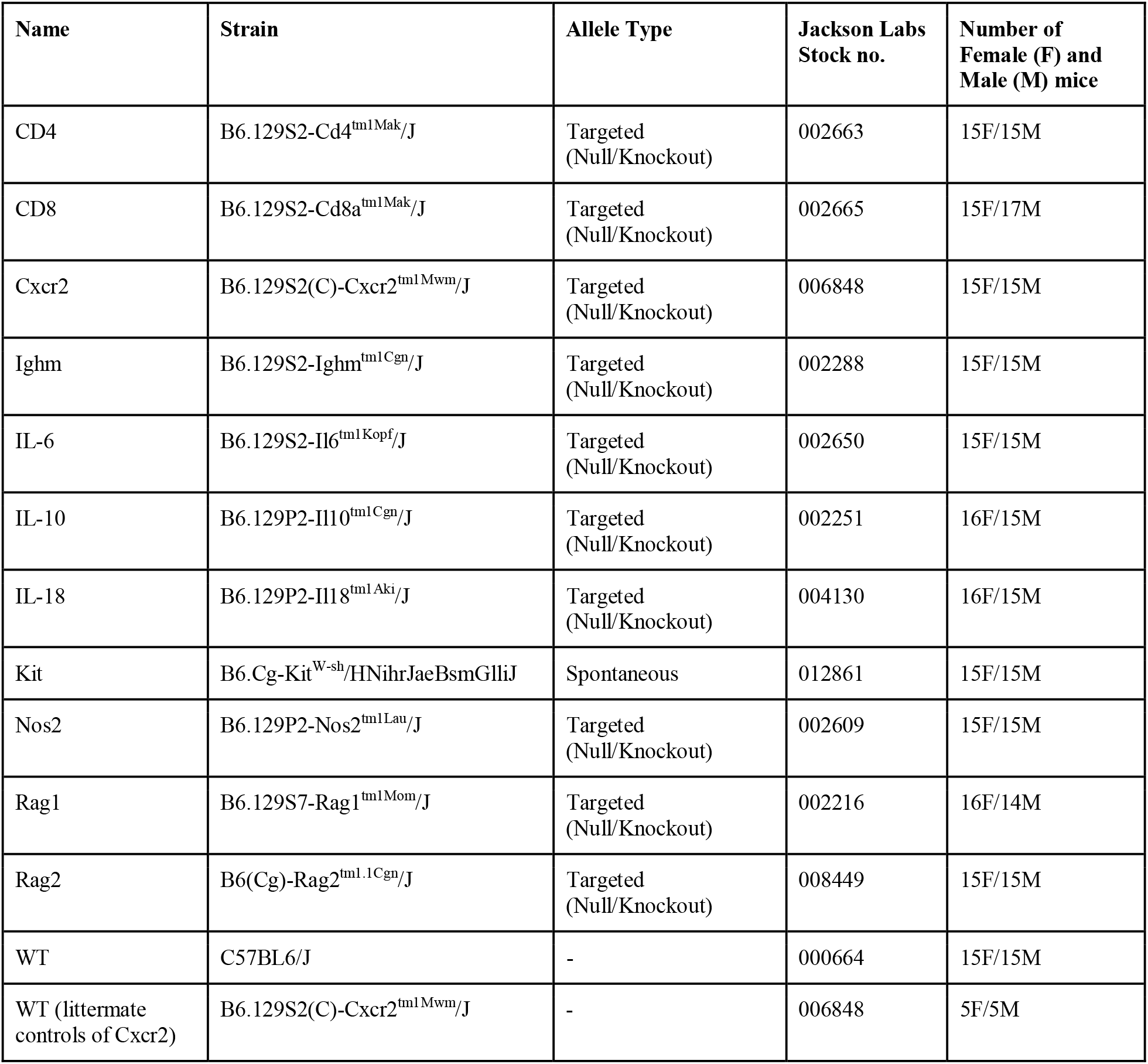
Information on strains used in this study.

Fixed brain samples were prepared for MRI using a previously described fixation protocol<u>[23]</u>. Briefly, mice were perfused with ProHance^®^ (gadoteridol, Bracco Diagnostics Inc., Princeton, NJ), and 4% paraformaldehyde (PFA) and then decapitated. Brains (kept within the skull) were separated from other tissue, allowed to postfix in a solution with PFA, ProHance^®^, and sodium azide (sodium trinitride, Fisher Scientific, Nepean, ON) at 4°C until imaging at 7-8 weeks postmortem<u>[24]</u>.

### Image acquisition and registration

A multi-channel 7.0-T MRI scanner with a 40 cm diameter bore (Varian Inc., Palo Alto, CA) was used to acquire 40-micron-isotropic anatomical images of 16 brains concurrently<u>[25]</u> in one 14-hour imaging session, with the following scan parameters: T2W 3D FSE cylindrical k-space acquisition sequence, TR/TE/ETL=350ms/12ms/6, two averages, FOV/matrix-size=20×20×25mm/504×504×630<u>[26]</u>. To quantify the anatomical differences between images, all 373 brain images were registered together using a previously described procedure<u>[27]</u>. Briefly, the mni_autoreg<u>[28]</u>, ANTS<u>[29]</u>, and pydpiper<u>[30]</u> toolkits were used to warp images to a consensus average. The MAGeT<u>[31]</u> pipeline was then used to automatically segment each of the 373 brain images into 336 structures using previously published segmentations<u>[32–36]</u> in order to determine structure volumes.

### Frequentist Statistics

All statistical analyses were conducted using the R software environment<u>[37–39]</u>. The two wild-type control strains were pooled for analyses as there were no significant differences between them. For each structure, volume was fit using a linear model with predictors of sex, strain, and their interactions; and partial F-test was used to assess significance. To determine direction of the strain effect, fitted coefficients were converted to t-statistics. Multiple comparisons were corrected using false-discovery-rate (FDR)<u>[40]</u>.

### Bayesian Statistics

Structure volumes were Z-transformed to standardize the volume data. An anatomical hierarchy<u>[41]</u> was used to reduce the number of structures (Supplementary Figure 3) by computing bayes-factors for the linear models’ residuals (examples shown in Supplementary Figure 4). Pruning the hierarchy (bayes-factor thresholded to 100) resulted in 95 bilateral brain structures.

A bayesian hierarchical model (BHM) was used to explore the effects of strain on neuroanatomical volumes of all 95 structures. It contained global and structure-specific predictors of sex, strain, and their interaction, and individual-specific intercept (priors given in Supplementary table 2). Stan<u>[42]</u> was used to fit the BHM (4 chains, 2500 sampling iterations each). The posterior distribution was used to compute the effect-size (*d*) marginalised over individual mice by subtracting the predicted volume (as a Z-score) of each mutant strain from the wild-type strain. For each structure, the probability of mutations resulting in large effect sizes (defined as |*d*|>1) was computed.

The posterior also provided the distribution of neuroanatomy phenotype as beta-values for each structure and strain (referenced to wild-type). Hellinger distance was used to assess dissimilarity of neuroanatomy phenotype between each pair of strains. Strains were annotated with anxiety phenotype based on existing literature (increased, decreased, or unchanged; summarised in Supplementary table 3). To assess whether strains with similar anxiety phenotype have lower Hellinger distances, permutation testing was performed (10^5^ iterations). Permutation tests were repeated excluding Cxcr2 (in which no anxiety phenotype has been reported in the literature) and including Cxcr2 on the assumption that the phenotype should be assigned ‘unchanged’, with similar results for both tests. To determine the effect of anxiety for each structure, the posterior was sampled 10^5^ times to generate volume distributions for all structures and mice. For each sample, a linear model (predictors of sex, anxiety, and their interaction) was fitted and anxiety effect size (*η*^2^) was calculated.

### Preferential gene expression and disease enrichment analysis

Using previously published methods<u>[43, 44]</u> and the Allen Brain Institute (ABI) gene expression atlas<u>[41]</u>, we explored the relationship between brain regions with altered volume and gene expression. Determined by the BHM, the top 25 brain structures with the highest absolute effect-size (i.e. all structures with |d|>1) constituted the region-of-interest (ROI) for this analysis. Preferential gene expression was evaluated using a fold-change measure: average gene expression in ROI divided by average gene expression in the brain.

Mouse genes were annotated with human diseases using two databases: the NCBI database<u>[45]</u> (mapping mouse genes into homologous humans genes) and DisGeNET database<u>[46]</u> (annotating human genes with associated diseases). For disease enrichment analysis, the target set was the top 4000 genes with the highest preferential expression and the background set was all homologous genes. Significance was assessed using hypergeometric tests with false-discovery-rate correction. Similar results were seen with target sets of the top 3000 and 5000 genes. When assessing significance of shifts in probability density, the Kolmogorov– Smirnov (KS) test was used. To investigate gene expression patterns through development, the ABI developmental gene expression atlas<u>[47]</u> was registered to the consensus average. Developmental gene expression data was clustered using k-means and the number of clusters determined using the elbow-method<u>[48]</u>.

## End Matter

### Notes

The authors declare no conflict of interest.

This article contains supporting information online.

## Acknowledgments

This work was supported by grants from the Canadian Institutes of Health Research as well as the Province of Ontario’s Neurodevelopmental Disorders (POND) network of the Ontario Brain Institute. Additional funding support was provided by the National Science and Engineering Research Council postgraduate scholarship (DF), the Hospital for Sick Children Restracomp Fellowship and Ontario Graduate Scholarship (DV).

## References

1. Estes ML, McAllister AK (2016) Maternal immune activation: Implications for neuropsychiatric disorders. Science 353:772–777

2. Knuesel I, Chicha L, Britschgi M, Schobel SA, Bodmer M, Hellings JA, Toovey S, Prinssen EP (2014) Maternal immune activation and abnormal brain development across CNS disorders. Nat Rev Neurol 10:643–660

3. Graves JS, Chitnis T, Weinstock-Guttman B, et al (2017) Maternal and Perinatal Exposures Are Associated With Risk for Pediatric-Onset Multiple Sclerosis. Pediatrics. https://doi.org/10.1542/peds.2016-2838

4. Atladóttir HO, Thorsen P, Østergaard L, Schendel DE, Lemcke S, Abdallah M, Parner ET (2010) Maternal infection requiring hospitalization during pregnancy and autism spectrum disorders. J Autism Dev Disord 40:1423–1430

5. Brown AS (2012) Epidemiologic studies of exposure to prenatal infection and risk of schizophrenia and autism. Dev Neurobiol 72:1272–1276

6. Smith SEP, Li J, Garbett K, Mirnics K, Patterson PH (2007) Maternal immune activation alters fetal brain development through interleukin-6. J Neurosci 27:10695–10702

7. Choi GB, Yim YS, Wong H, Kim S, Kim H, Kim SV, Hoeffer CA, Littman DR, Huh JR (2016) The maternal interleukin-17a pathway in mice promotes autism-like phenotypes in offspring. Science 351:933–939

8. Ponzio NM, Servatius R, Beck K, Marzouk A, Kreider T (2007) Cytokine levels during pregnancy influence immunological profiles and neurobehavioral patterns of the offspring. Ann N Y Acad Sci 1107:118–128

9. Chourbaji S, Urani A, Inta I, Sanchis-Segura C, Brandwein C, Zink M, Schwaninger M, Gass P (2006) IL-6 knockout mice exhibit resistance to stress-induced development of depression-like behaviors. Neurobiol Dis 23:587–594

10. Mesquita AR, Correia-Neves M, Roque S, Castro AG, Vieira P, Pedrosa J, Palha JA, Sousa N (2008) IL-10 modulates depressive-like behavior. J Psychiatr Res 43:89–97

11. Yaguchi T, Nagata T, Yang D, Nishizaki T (2010) Interleukin-18 regulates motor activity, anxiety and spatial learning without affecting synaptic plasticity. Behav Brain Res 206:47–51

12. Rilett KC, Friedel M, Ellegood J, MacKenzie RN, Lerch JP, Foster JA (2015) Loss of T cells influences sex differences in behavior and brain structure. Brain Behav Immun 46:249–260

13. Clark SM, Soroka JA, Song C, Li X, Tonelli LH (2016) CD4(+) T cells confer anxiolytic and antidepressant-like effects, but enhance fear memory processes in Rag2(-/-) mice. Stress 19:303–311

14. Sankar A, MacKenzie RN, Foster JA (2012) Loss of class I MHC function alters behavior and stress reactivity. J Neuroimmunol 244:8–15

15. Rattazzi L, Piras G, Ono M, Deacon R, Pariante CM, D’Acquisto F (2013) CD4^+^ but not CD8^+^ T cells revert the impaired emotional behavior of immunocompromised RAG-1-deficient mice. Transl Psychiatry 3:e280

16. Clark SM, Sand J, Francis TC, Nagaraju A, Michael KC, Keegan AD, Kusnecov A, Gould TD, Tonelli LH (2014) Immune status influences fear and anxiety responses in mice after acute stress exposure. Brain Behav Immun 38:192–201

17. Buskila Y, Abu-Ghanem Y, Levi Y, Moran A, Grauer E, Amitai Y (2007) Enhanced astrocytic nitric oxide production and neuronal modifications in the neocortex of a NOS2 mutant mouse. PLoS One 2:e843

18. Nautiyal KM, Ribeiro AC, Pfaff DW, Silver R (2008) Brain mast cells link the immune system to anxiety-like behavior. Proc Natl Acad Sci U S A 105:18053–18057

19. Ellegood J, Anagnostou E, Babineau BA, et al (2015) Clustering autism: using neuroanatomical differences in 26 mouse models to gain insight into the heterogeneity. Mol Psychiatry 20:118–125

20. Anacker C, Scholz J, O’Donnell KJ, Allemang-Grand R, Diorio J, Bagot RC, Nestler EJ, Hen R, Lerch JP, Meaney MJ (2016) Neuroanatomic Differences Associated With Stress Susceptibility and Resilience. Biol Psychiatry 79:840–849

21. Cacalano G, Lee J, Kikly K, Ryan AM, Pitts-Meek S, Hultgren B, Wood WI, Moore MW (1994) Neutrophil and B cell expansion in mice that lack the murine IL-8 receptor homolog. Science 265:682–684

22. Holmdahl R, Malissen B (2012) The need for littermate controls. Eur J Immunol 42:45–47

23. Cahill LS, Laliberté CL, Ellegood J, Spring S, Gleave JA, van Eede MC, Lerch JP, Henkelman RM (2012) Preparation of fixed mouse brains for MRI. Neuroimage 60:933–939

24. de Guzman AE, de Guzman AE, Wong MD, Gleave JA, Nieman BJ (2016) Variations in post-perfusion immersion fixation and storage alter MRI measurements of mouse brain morphometry. NeuroImage 142:687–695

25. Dazai J, Spring S, Cahill LS, Henkelman RM (2011) Multiple-mouse neuroanatomical magnetic resonance imaging. J Vis Exp. https://doi.org/10.3791/2497

26. Spencer Noakes TL, Henkelman RM, Nieman BJ (2017) Partitioning k-space for cylindrical three-dimensional rapid acquisition with relaxation enhancement imaging in the mouse brain. NMR Biomed. https://doi.org/10.1002/nbm.3802

27. Lerch JP, Yiu AP, Martinez-Canabal A, Pekar T, Bohbot VD, Frankland PW, Henkelman RM, Josselyn SA, Sled JG (2011) Maze training in mice induces MRI-detectable brain shape changes specific to the type of learning. Neuroimage 54:2086–2095

28. Collins DL, Neelin P, Peters TM, Evans AC (1994) Automatic 3D intersubject registration of MR volumetric data in standardized Talairach space. J Comput Assist Tomogr 18:192–205

29. Avants BB, Tustison NJ, Song G, Cook PA, Klein A, Gee JC (2011) A reproducible evaluation of ANTs similarity metric performance in brain image registration. Neuroimage 54:2033–2044

30. Friedel M, van Eede MC, Pipitone J, Chakravarty MM, Lerch JP (2014) Pydpiper: a flexible toolkit for constructing novel registration pipelines. Front Neuroinform 8:67

31. Chakravarty MM, Steadman P, van Eede MC, Calcott RD, Gu V, Shaw P, Raznahan A, Collins DL, Lerch JP (2013) Performing label-fusion-based segmentation using multiple automatically generated templates. Hum Brain Mapp 34:2635–2654

32. Dorr AE, Lerch JP, Spring S, Kabani N, Henkelman RM (2008) High resolution three-dimensional brain atlas using an average magnetic resonance image of 40 adult C57Bl/6J mice. Neuroimage 42:60–69

33. Steadman PE, Ellegood J, Szulc KU, Turnbull DH, Joyner AL, Henkelman RM, Lerch JP (2014) Genetic effects on cerebellar structure across mouse models of autism using a magnetic resonance imaging atlas. Autism Res 7:124–137

34. Ullmann JFP, Watson C, Janke AL, Kurniawan ND, Reutens DC (2013) A segmentation protocol and MRI atlas of the C57BL/6J mouse neocortex. Neuroimage 78:196–203

35. Richards K, Watson C, Buckley RF, et al (2011) Segmentation of the mouse hippocampal formation in magnetic resonance images. Neuroimage 58:732–740

36. Qiu LR, Fernandes DJ, Szulc-Lerch KU, Dazai J, Nieman BJ, Turnbull DH, Foster JA, Palmert MR, Lerch JP (2018) Mouse MRI shows brain areas relatively larger in males emerge before those larger in females. Nat Commun 9:2615

37. (2010) R: A Language and Environment for Statistical Computing : Reference Index.

38. Wickham H, Grolemund G (2016) R for Data Science: Import, Tidy, Transform, Visualize, and Model Data. “O’Reilly Media, Inc.”

39. Nalborczyk L, Batailler C, Lœvenbruck H, Vilain A, Bürkner P-C (2019) An Introduction to Bayesian Multilevel Models Using brms: A Case Study of Gender Effects on Vowel Variability in Standard Indonesian. J Speech Lang Hear Res 62:1225–1242

40. Genovese CR, Lazar NA, Nichols T (2002) Thresholding of statistical maps in functional neuroimaging using the false discovery rate. Neuroimage 15:870–878

41. Lein ES, Hawrylycz MJ, Ao N, et al (2007) Genome-wide atlas of gene expression in the adult mouse brain. Nature 445:168–176

42. Korner-Nievergelt F, Roth T, von Felten S, Guélat J, Almasi B, Korner-Nievergelt P (2015) Bayesian Data Analysis in Ecology Using Linear Models with R, BUGS, and Stan. Academic Press

43. Fernandes DJ, Ellegood J, Askalan R, et al (2017) Spatial gene expression analysis of neuroanatomical differences in mouse models. Neuroimage 163:220–230

44. Fernandes DJ, Spring S, Roy AR, Qiu LR, Yee Y, Nieman BJ, Lerch JP, Palmert MR (2021) Exposure to maternal high-fat diet induces extensive changes in the brain of adult offspring. Transl Psychiatry 11:149

45. Wheeler DL (2004) Database resources of the National Center for Biotechnology Information. Nucleic Acids Research 33:D39–D45

46. Piñero J, Queralt-Rosinach N, Bravo À, Deu-Pons J, Bauer-Mehren A, Baron M, Sanz F, Furlong LI (2015) DisGeNET: a discovery platform for the dynamical exploration of human diseases and their genes. Database 2015:bav028

47. Thompson CL, Ng L, Menon V, et al (2014) A high-resolution spatiotemporal atlas of gene expression of the developing mouse brain. Neuron 83:309–323

48. Thorndike RL (1953) Who belongs in the family? Psychometrika 18:267–276

49. Wootla B, Eriguchi M, Rodriguez M (2012) Is multiple sclerosis an autoimmune disease? Autoimmune Dis 2012:969657

50. Wasielewska JM, Grönnert L, Rund N, Donix L, Rust R, Sykes AM, Hoppe A, Roers A, Kempermann G, Walker TL (2017) Mast cells increase adult neural precursor proliferation and differentiation but this potential is not realized in vivo under physiological conditions. Sci Rep 7:17859

51. Calhoon GG, Tye KM (2015) Resolving the neural circuits of anxiety. Nat Neurosci 18:1394–1404

52. Apps R, Strata P (2015) Neuronal circuits for fear and anxiety - the missing link. Nat Rev Neurosci 16:642

53. Anthony TE, Dee N, Bernard A, Lerchner W, Heintz N, Anderson DJ (2014) Control of stress-induced persistent anxiety by an extra-amygdala septohypothalamic circuit. Cell 156:522–536

54. Moreno-Rius J (2018) The cerebellum in fear and anxiety-related disorders. Prog Neuropsychopharmacol Biol Psychiatry 85:23–32

55. Koutsikou S, Crook JJ, Earl EV, Leith JL, Watson TC, Lumb BM, Apps R (2014) Neural substrates underlying fear-evoked freezing: the periaqueductal grey-cerebellar link. J Physiol 592:2197–2213

56. Silveira C, Guedes R, Maia D, Curral R, Coelho R (2019) Neuropsychiatric Symptoms of Multiple Sclerosis: State of the Art. Psychiatry Investig 16:877–888

57. Broen MPG, Narayen NE, Kuijf ML, Dissanayaka NNW, Leentjens AFG (2016) Prevalence of anxiety in Parkinson’s disease: A systematic review and meta-analysis. Mov Disord 31:1125–1133

58. Didonna A, Oksenberg JR (2017) The Genetics of Multiple Sclerosis. Multiple Sclerosis: Perspectives in Treatment and Pathogenesis

59. Mesaros S, Rocca MA, Riccitelli G, et al (2009) Corpus callosum damage and cognitive dysfunction in benign MS. Hum Brain Mapp 30:2656–2666

60. Harrison DM, Oh J, Roy S, et al (2015) Thalamic lesions in multiple sclerosis by 7T MRI: Clinical implications and relationship to cortical pathology. Mult Scler 21:1139–1150

61. Bermel RA, Innus MD, Tjoa CW, Bakshi R (2003) Selective caudate atrophy in multiple sclerosis: a 3D MRI parcellation study. Neuroreport 14:335–339

62. Quint DJ, Cornblath WT, Trobe JD (1993) Multiple sclerosis presenting as Parinaud syndrome. AJNR Am J Neuroradiol 14:1200–1202

63. Jiang T, Li G, Xu J, Gao S, Chen X (2018) The Challenge of the Pathogenesis of Parkinson’s Disease: Is Autoimmunity the Culprit? Front Immunol 9:2047

64. Jevtic S, Sengar AS, Salter MW, McLaurin J (2017) The role of the immune system in Alzheimer disease: Etiology and treatment. Ageing Res Rev 40:84–94

