## Supplementary Figures for "Mouse models of immune dysfunction: Their neuroanatomical differences reflect their anxiety-behavioural phenotype"

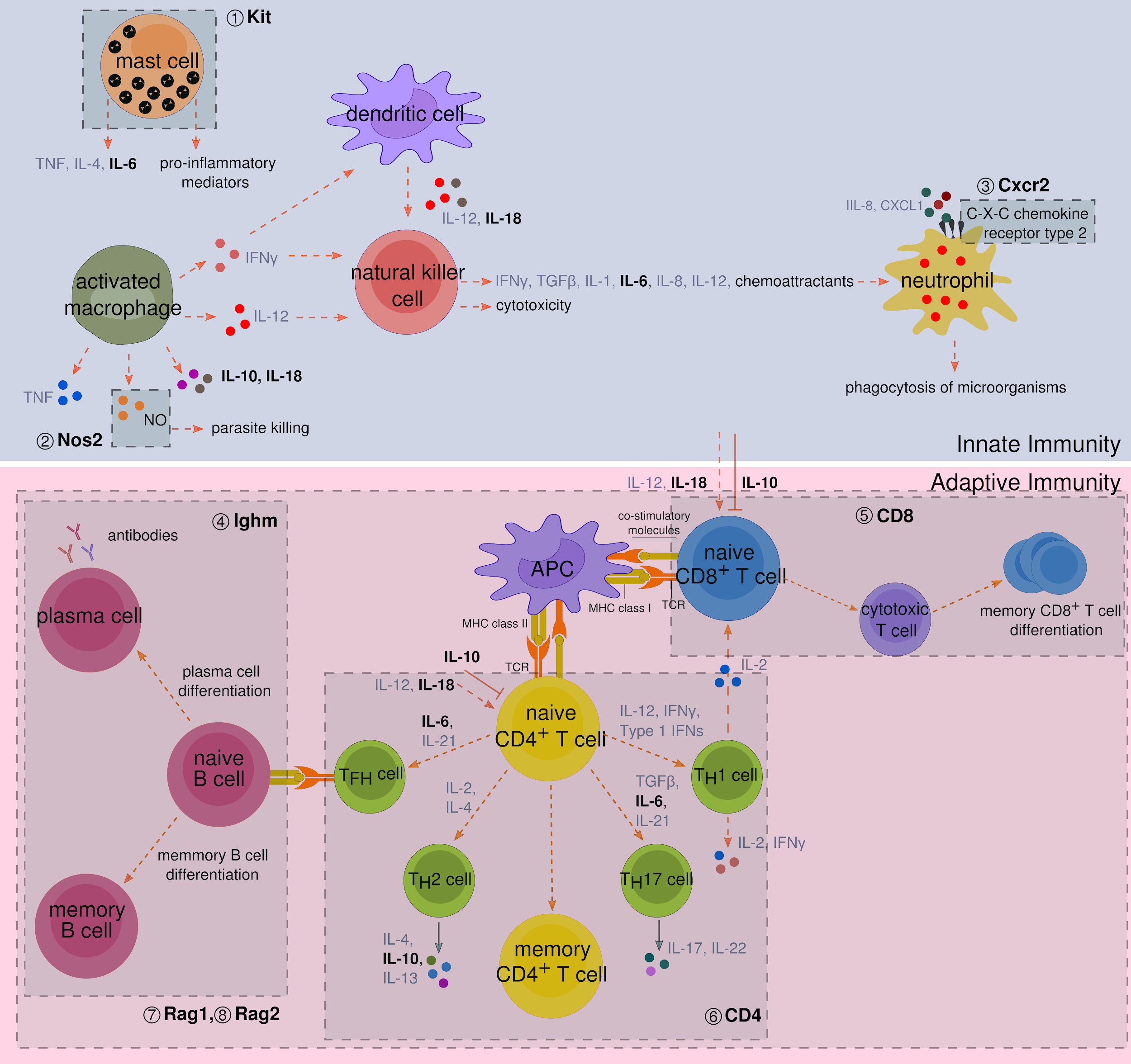

**Supplementary Figure 1:** Schematic representation depicting the functions of the various immune system components mutated in this study. Kit ① mice are depleted of mast cells soon after birth, affecting inflammatory responses and cytokine release. Nos2 ② mutants lack serum nitric oxide responses involved in parasite killing. Cxcr2 ③ mutants lack the chemokine receptor that binds CXCL2 and IL-8 leading to impaired neutrophil recruitment and decreased pathogen clearance. The Ighm ④ mutant lacks mature B cells impeding humoral immunity. CD8 ⑤ knockouts are deficient in functional cytotoxic T cells while CD4 ⑥ mutants have a significant block in CD4+ T cell development. Both Rag1 ⑦ and Rag2 ⑧ mutants lack mature, functional B and T cells and are therefore deficient in adaptive immune responses. The cytokines studied in this paper -- IL-6, IL-10 and IL-18 -- are shown in bold and act on multiple pathways. IL-6 is both a pro-inflammatory cytokine and anti-inflammatory myokine while IL-10 is an anti-inflammatory cytokine. IL-18 is a pro-inflammatory cytokine playing a key role in autoimmune, inflammatory and infectious diseases. This figure is not meant to be a comprehensive depiction of the immune system and all components. APC, antigen presenting cell; MHC, major histocompatibility complex, TCR, T cell receptor, T_H_, T helper cell, T_FH_, T follicular helper cell.

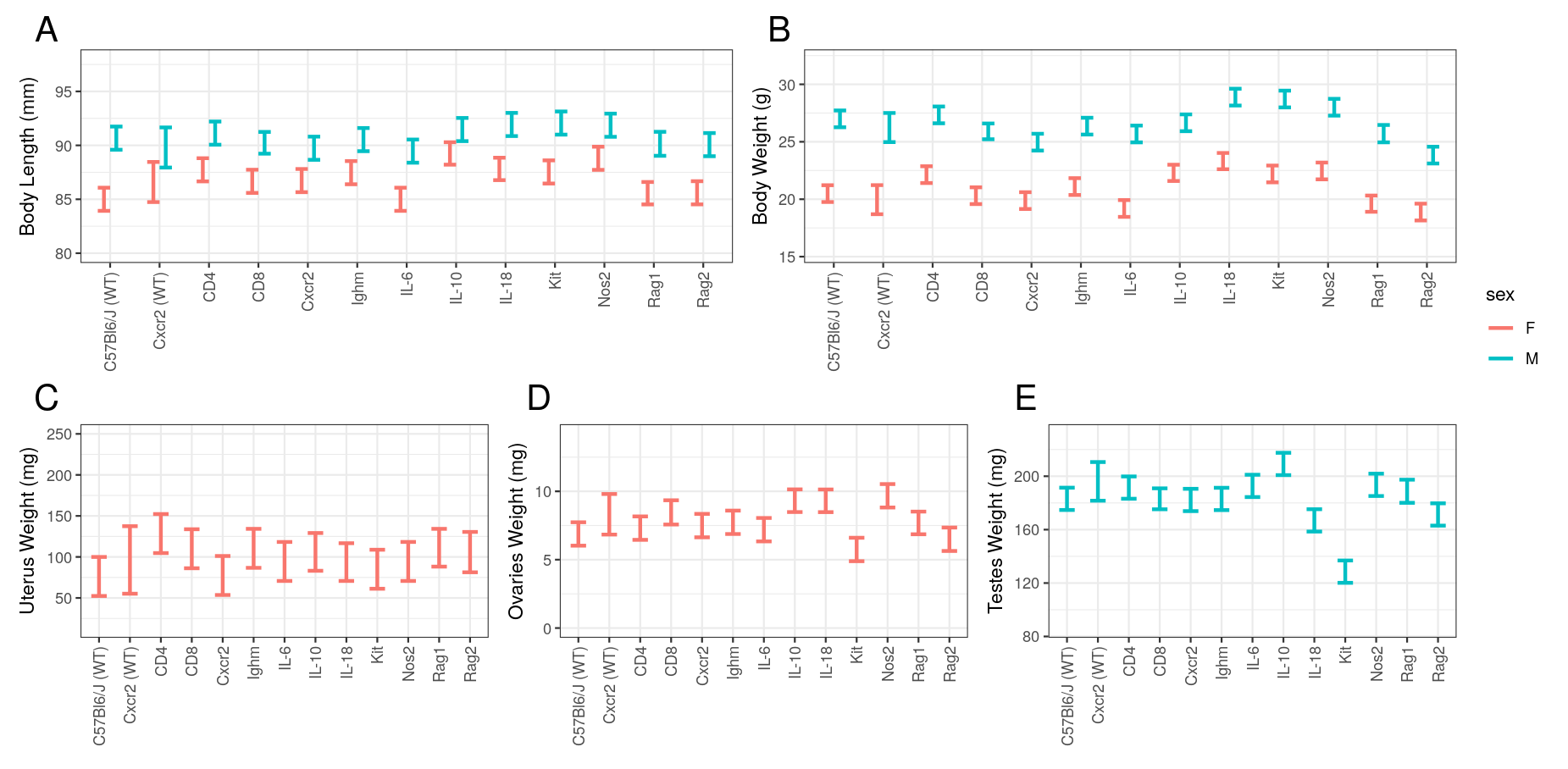

**Supplementary Figure 2:** Physical characteristics of each strain in study. (A) Body length, (B) body weight, (C) uterus weight, (D) ovaries weight, and (E) testes weight were measured. There was a significant effect of strain on body length (F_24,372_ = 4.5,p < 10^-10^), body weight (F_24,372_ = 14.9,p < 10^-40^), ovaries weight (F_12,186_ = 7.2,p < 10^-9^), and testes weight (F_12,185_ = 20.4,p < 10^-27^), but not uterus weight (F_12,186_ = 1.5,p = 0.12). There was no significant sex-strain interactions in the body length (F_12,360_ = 1.43,p = 0.15) and body weight (F_12,360_ = 1.63,p = 0.08) measures.

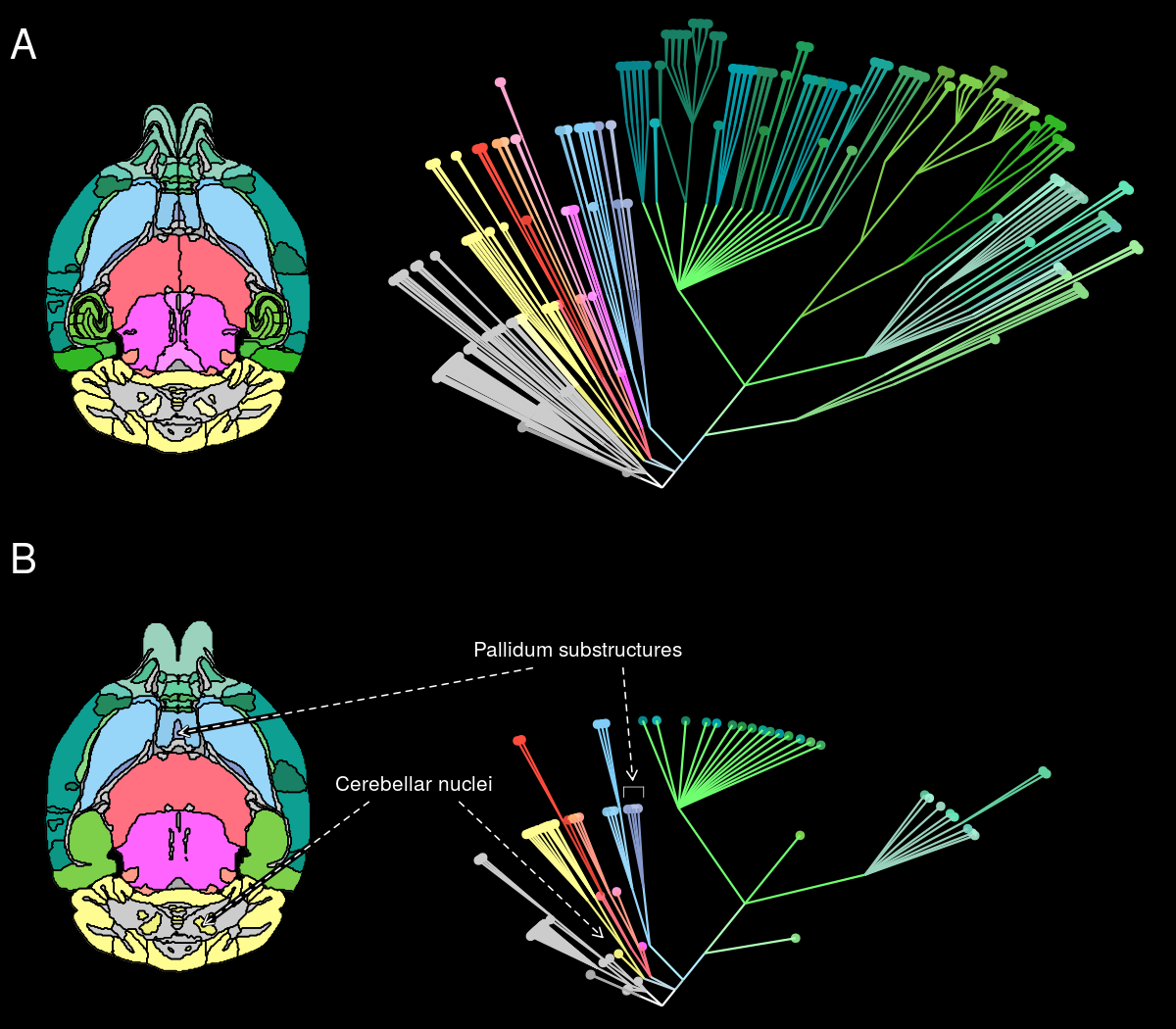

**Supplementary Figure 3:** Pruning the neuroanatomy hierarchy. (A) All 336 structures in the mouse brain atlas (left) can be organised into a developmental hierarchy (right). The first level of the hierarchy merges bilateral structures. In all subsequent levels, substructures merge together to form larger structures. The root of the tree represents the whole brain. (B) Pruning the hierarchy reduces the number of total structures by merging together substructures that have similar phenotypes across sex and strain. For example, the substructures of the cerebellar nuclei have similar phenotypes and thus they are merged together after pruning the tree. On the other hand, the pallidum substructures do not have similar phenotype and are, therefore, not pruned. Volume data for these examples are shown in Supplementary Figure 4.

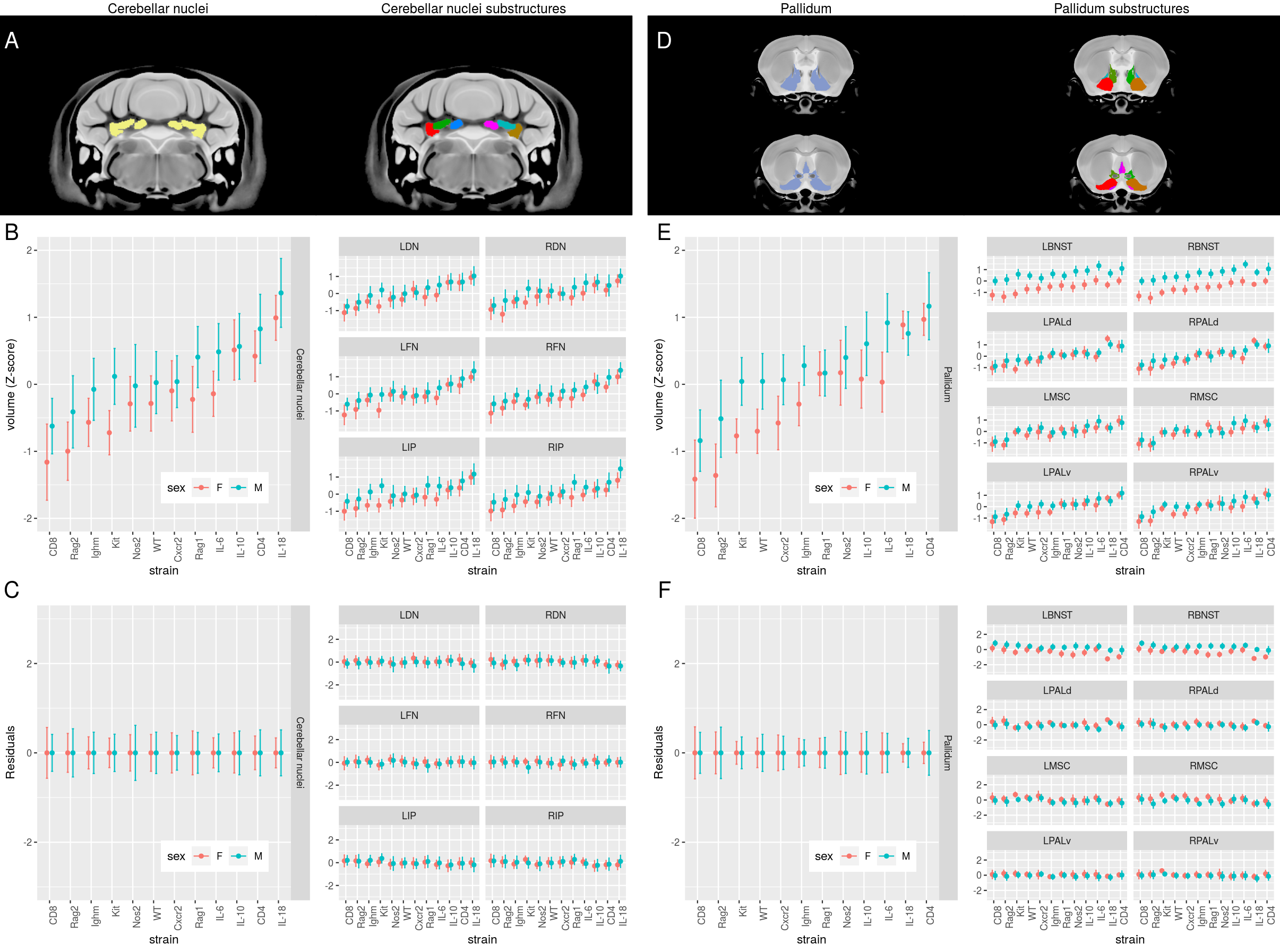

**Supplementary Figure 4:** Illustration of pruning neuroanatomy hierarchy. (A) Segmentation of Cerebellar nuclei (Left) and its substructures (Right): Left Dentate Nucleus (LDN), Right Dentate Nucleus (RDN), Left Fastigial Nucleus (LFN), Right Fastigial Nucleus (LFN), Left Interposed Nucleus (LIP), and Right Interposed Nucleus (RIP). (B) Volume (as a Z-score) for the Cerebellar nuclei and each substructure. (C) Residuals from the model fitted to Cerebellar nuclei. Substructure residuals are not influenced by sex and strain, indicating that substructure response to sex and strain is well-accounted by the model fitted to the parent structure. This can be quantified rigorously using bayes factor comparing two models fitted to substructure residuals: intercept-only model vs sex and strain (with interaction) model. The intercept-only model is highly favoured (bayes factor for all substructures exceeded 10^18^). (D) Segmentation of Pallidum (Left) and its substructures (Right): Left Bed Nucleus of the Stria Terminalis (LBNST), Right Bed Nucleus of the Stria Terminalis (RBNST), Left Dorsal Pallidum (LPALd), Right Dorsal Pallidum (RPALd), Left Medial Septal Complex (LMSC), Right Medial Septal Complex (RMSC), Left Ventral Pallidum (LPALv), and Right Ventral Pallidum (RPALv). (E) Volume (as a Z-score) for the Pallidum and substructures. (F) Residuals from the model fitted to Pallidum. Substructure residuals are influenced by sex and strain, indicating that substructure response to sex and strain is not well-accounted by the model fitted to the parent structure: Bayes factor comparing two models fitted to substructures -- intercept-only model vs sex and strain (with interaction) model -- showed this to be the case: intercept-only model is highly unfavourable (bayes factor for RBNST is <10^-12^).

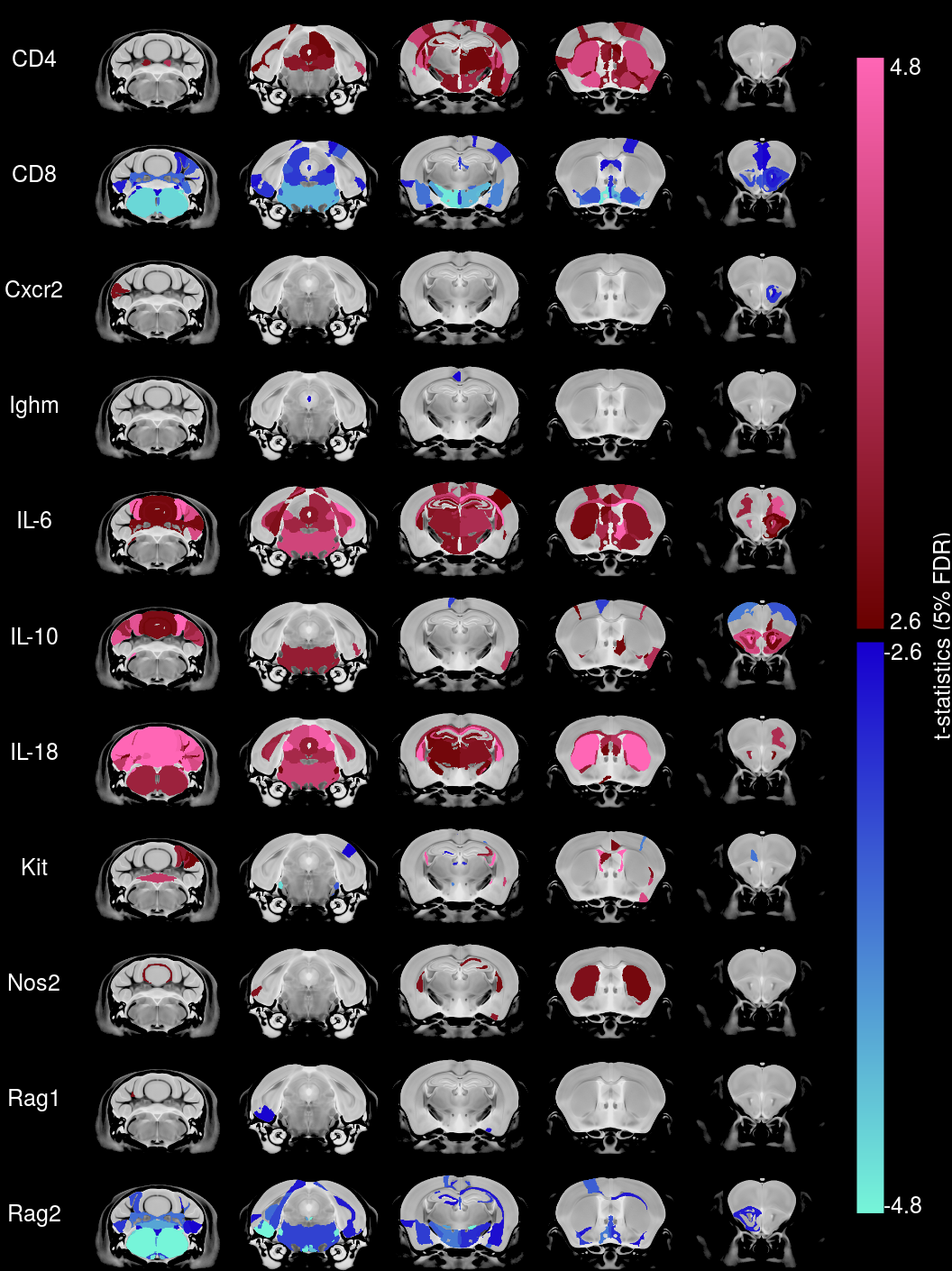

**Supplementary Figure 5:** Immune system mutations have a highly heterogeneous effect on mouse brain anatomy. The directional effect in males of the various mutant strains relative to the wild-type strains is visualized using t-statistics and shows a heterogeneous neuroanatomical phenotype. Regions larger or smaller in mutants relative to wild-type are given maroon-pink and blue-turquoise colours, respectively, if effects are <5%FDR. Saturated colours represent effects <0.01% FDR.

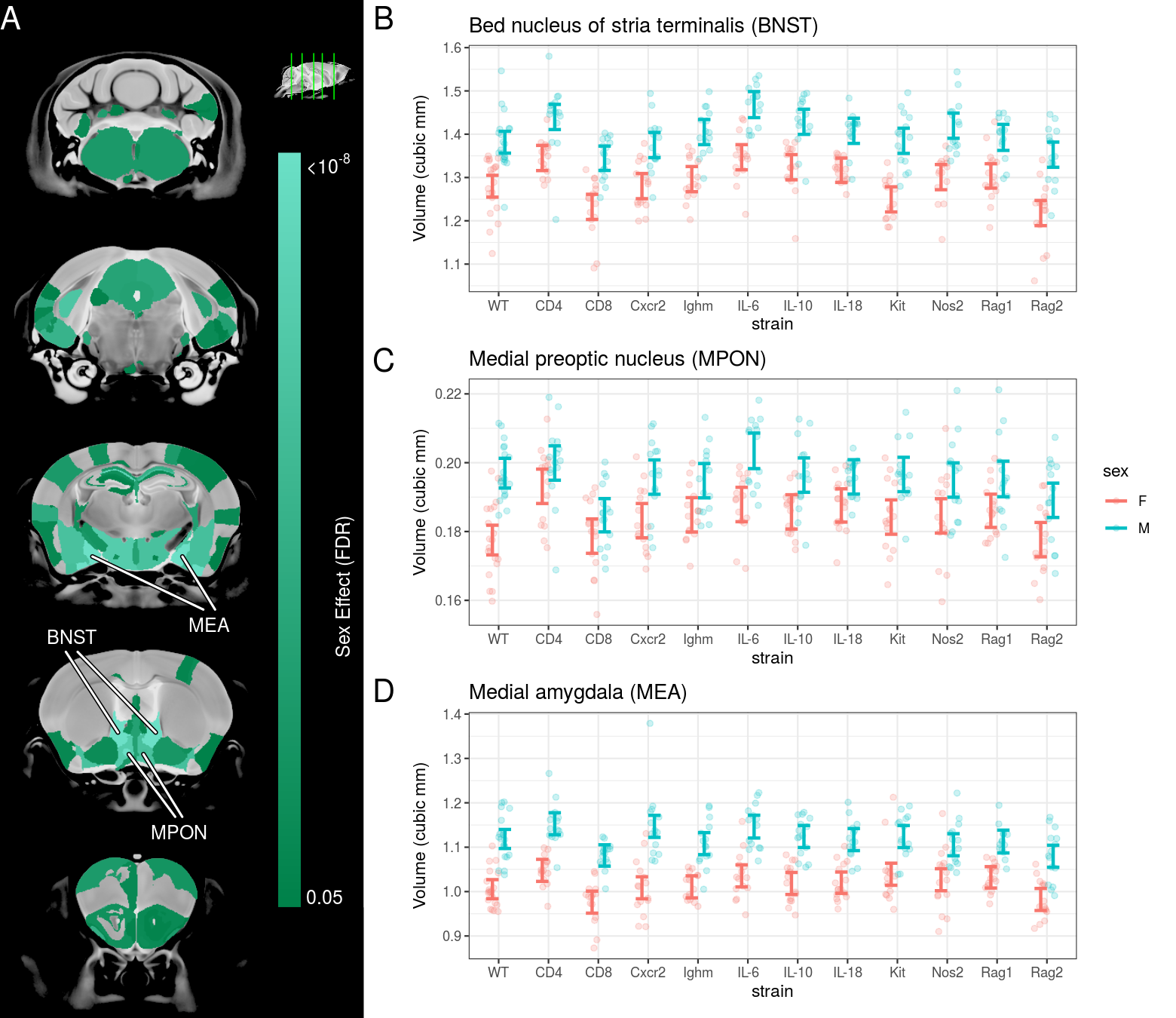

**Supplementary Figure 6:** Immune system mutations have a significant effect on the neuroanatomy of both sexes. (A) Sexual dimorphisms were also found throughout the brain, however no significant interactions between sex and the immune strains were identified. (B-D) The volume of canonical sexually dimorphic structures -- Bed nucleus of stria terminalis (BNST), Medial preoptic nucleus (MPON), Medial amygdala (MEA) -- in each strain are shown. These structures showed a significant effect of strain (BNST, F_22,369_=6.87,p<10^-16^; MPON, F_22,369_=3.49,p<10^-6^; MEA, F_22,369_=3.24,p<10^-5^) and sex (BNST, F_12,359_=28.37,p<10^-43^; MPON, F_12,359_=11.29,p<10^-19^; MEA, F_12,359_=34.05, p<10^-50^), but no significant sex-strain interaction (BNST, F_11,358_=0.59,p=0.8; MPON, F_11,358_=1.23,p=0.26; MEA, F_11,358_=0.84,p=0.6). Colour maps show effects under 5% FDR and error bars represent 95% confidence intervals.
